# Degradation determinants are abundant in human noncanonical proteins

**DOI:** 10.1101/2024.05.01.592071

**Authors:** Claudio Casola, Adekola Owoyemi, Nikolaos Vakirlis

**Author notes:** Corresponding Authors Information: Claudio Casola: 534 John Kimbrough Blvd, TAMU 2258, College Station, TX 77843-2258, Nikolaos Vakirlis: 34 Fleming Street 16672, 16672 Vari, Greece.

## Abstract

The comprehensive characterization of human proteins, a key objective in contemporary biology, has been revolutionized by the identification of thousands of potential novel proteins through ribosome profiling and proteomics. Determining the physiological activity of these noncanonical proteins has proven difficult, because they are encoded by different types of coding regions and tend to share no sequence similarity with canonical polypeptides. Evidence from immunopeptidomic assays combined with a better understanding of the quality control of protein synthesis suggest that many noncanonical proteins may possess low stability in the cellular environment. Here, we tested this hypothesis by analyzing the frequency of multiple sequence features eliciting either proteasomal degradation or autophagy across 91,003 canonical (annotated) protein isoforms and 232,460 noncanonical proteins. Overall, noncanonical proteins were enriched for degradation-related features compared to all canonical proteins. Notably, degradation determinants were also enriched in canonical protein isoforms starting with a non-methionine amino acid. Analyses of original and shuffled sequences showed evidence of selective pressure either against or towards the accumulation of specific degradation signatures only in major isoforms of canonical proteins. However, stability was significantly higher in noncanonical proteins with evidence of phenotypic effects when knocked-out in cell lines. Notably, we found that the C-terminal tail hydrophobicity represents a reliable proxy for degradation propensity with potential applications in identifying functional noncanonical proteins. These findings underscore the critical role of degradation processes in regulating the half-life of noncanonical proteins and demonstrate the power of degradation-associated signatures in discriminating noncanonical genes likely to encode for biologically functional molecules.

## Introduction

Annotating all proteins encoded by the human genome is paramount to our understanding of human physiology, disease and evolution. The number of known, or ‘canonical’, protein coding genes has been shown to be around 20,000, with many genes encoding multiple protein isoforms derived from alternative splicing (1). However, it is becoming increasingly appreciated that human genome harbors many other noncanonical open reading frames (ncORFs) with the potential to encode polypeptides (2–4). These noncanonical ORFs disproportionally encode for small proteins that have largely been overlooked by gene annotation algorithms and are sometimes referred to as the ‘dark proteome’ (5, 6).

The development of ribosome profiling (Ribo-seq) technologies has dramatically increased the ability to identify potentially translated ncORFs (7–16). Additionally, proteomic data have directly showed synthesis of noncanonical proteins (14, 17–19), whereas evolutionary analyses revealed selective constraints compatible with protein-coding sequences in a subset of ncORFs (8, 20). Nevertheless, a physiological role has been conclusively demonstrated only for a limited number of noncanonical proteins, or NCPs, raising the question of how many of these sequences constitute functional polypeptides. The uncertainties surrounding the proportion of bioactive NCPs are also apparent from annotation efforts, with estimates of these proteins spanning three orders of magnitude (21–23).

Some proteomic evidence suggests that many NCPs might represent aberrant translation products that are rapidly degraded. For instance, NCPs tend to be disproportionally represented in the immunopeptidome—the collection of peptides from proteasomal cleavage that are presented by the MHC I—compared to the whole-proteasome (11–13, 24). This evidence and other observations have prompted the development of the “defective ribosomal products” (DRiPs) hypothesis, which postulates that a large proportion of NCPs represent misfolded polypeptides or translation errors that are degraded during or soon after translation, feeding the resulting peptides into the immunopeptidome (25, 26).

The DRiPs hypothesis has received further support by recent analyses revealing that NCPs on average possess higher hydrophobicity at the C-terminal tail than canonical proteins (hereafter, CPs), a feature associated with proteasomal proteolysis (27). Additionally, several types of ncORFs detected via Ribo-seq might not result in a protein product. Indeed, some upstream ORFs (uORFs) regulate translation of the main ORF on the same transcript by interacting with ribosomes without producing a protein (28). Similarly, translation of lncRNA transcripts based on Ribo-seq data alone has been questioned (29, 30).

Nevertheless, there is increasing support for the view that many human noncanonical proteins are functional. Knock-out experiments have shown that thousands of NCPs exhibit a phenotypic effect, albeit in cell cultures (10), and a significant proportion of NCPs have been detected across multiple proteomic sequencing data (22). Moreover, the depletion of NCPs in whole-proteome datasets could be attributable to the different distribution of tryptic sites between canonical and noncanonical proteins, resulting in a high rate of NCP false negatives (31, 32). Experimental analyses integrating Ribo-seq and mass-spectrometry data can further improve detection of stable NCPs (31); however, only a few such studies have been developed so far, with only a subset focusing specifically on NCPs (11, 12).

Computational predictions of sequence signatures associated with protein stability can significantly contribute to determining the functional impact of NCPs, both complementing and expanding on translation surveys and on the limited number of well-characterized noncanonical proteins. For instance, stability profiles of NCPs can inform on sequences that are less likely to be rapidly degraded and therefore should be prioritized for in-depth experimental analyses.

In this study, we investigated the stability of noncanonical proteins using an array of experimentally characterized protein degradation determinants linked with either proteasomal digestion or autophagy. We analyzed 91,003 canonical proteins and 232,460 putative NCPs from seven Ribo-seq datasets and meta datasets on multiple human cell lines and tissues. Our results showed a significant enrichment of all degradation predictors in noncanonical proteins. Notably, these signatures were less frequent in some NCP categories that are more likely to be functional, including those with a reported phenotypic effect following knock-out experiments. Our findings allowed the identification of promising functional noncanonical protein candidates and inform the ongoing annotation efforts of the human genome and proteome.

## Results

### Datasets and degradation determinants

Human noncanonical proteins have been obtained from a variety of tissues and cell types throughout several methodological approaches. In order to determine if noncanonical proteins show consistent patterns of degradation predictors we analyzed data from 5 individual ribosomal profiling datasets and two meta-datasets (9, 10, 16, 21, 22, 33, 34), in addition to protein sequences encoded by ∼20,000 canonical genes (Table S1). For each sequence, we inferred four degradation determinants: (1) The combined hydrophobicity of the last 30 amino acids of each protein sequence, or C-terminal tail hydrophobicity scores (hereafter CTTH); (2) The terminal (N-end and C-end) degrons; (3) The terminal intrinsic disorder regions, or IDRs; and (4) The microautophagy-associated KFERQ-like motifs (Fig. 1A).

**Figure 1.**
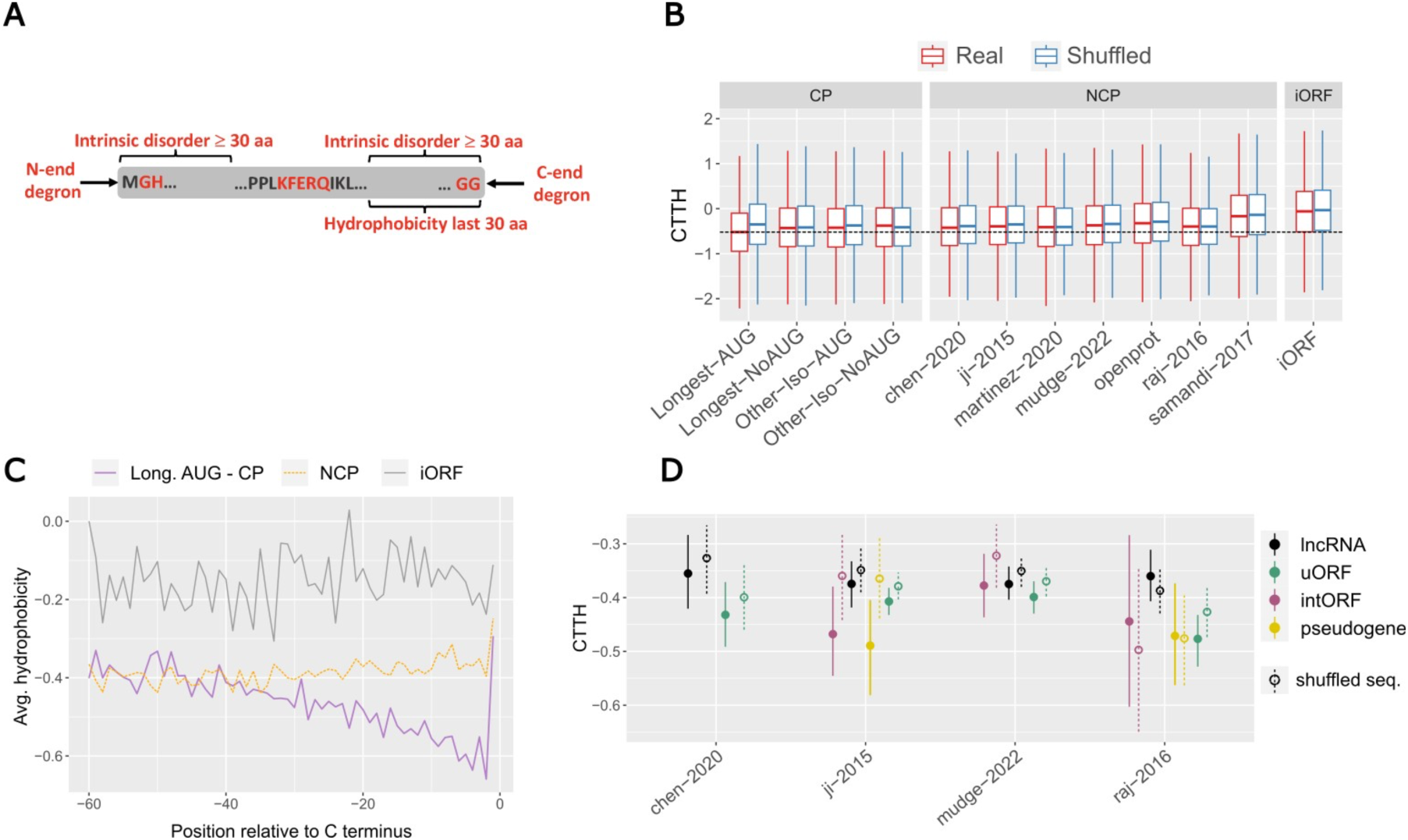
**A.** Representation of the four degradation determinants analyzed in this study and their localization along protein sequences along a protein sequence (gray rectangle). The N-end and C-end degron motifs correspond to two recently reported strong degrons [42-44] A KFERQ-like motif is shown around the center of the protein. **B.** C-terminal tail hydrophobicity scores (CTTH) calculated using the Kyte-Doolittle scale in AUG and non-AUG CPs and in NCP datasets. Values for both original (red) and shuffled (blue) sequences are shown. iORFs: nontranslated intergenic unannotated ORFs. Black dotted line represents the average for Longest-AUG dataset. **C.** Average hydrophobicity (Kyte-Doolittle scale) in the last 60 amino acids of CPs, NCPs and iORFs. **D.** CTTH in noncanonical protein types, in real and shuffled sequences.

### C-terminal tail hydrophobicity

It has recently been shown that an elevated average hydrophobicity in the last 30 amino acids of protein sequence is strongly associated with protein degradation via the proteasome-ubiquitin system (27) (Fig. 1A). We calculated this parameter, known as C-terminal tail hydrophobicity scores (CTTH), across all CPs and NCPs using the Kyte-Doolittle scale (35), in which hydrophobicity corresponds to positive values. Although CTTH was on average negative (hydrophilic) in all datasets, we observed significant variation in the overall hydropathy. The longest canonical isoforms showed the lowest average CTTH (−0.52), indicating diminished hydrophobicity, compared to other canonical isoforms (Wilcoxon rank sum test, see Methods, *P* < 5.47 x 10^-8^ in all tests, Fig. 1B, Table S2). Moreover, the CTTH score was on average lower and less variable in canonical isoforms starting with a methionine, or AUG isoforms, compared to non-AUG isoforms (Fig. 1B).

Across NCP datasets, CTTH was significantly higher than the longest-AUG canonical isoforms, ranging between −0.17 and −0.41 (*P* < 2.99 x 10^-5^ in all tests, Fig. 1B, Table S2). The Samandi-2017 (16) dataset includes a large group of ORFs with limited evidence of translation, possibly explaining the much higher CTTH. Putative proteins encoded by non-translated intergenic ORFs (iORF proteins) showed the highest CTTH score. Overall, protein size explained little to no variance in CTTH scores—all datasets had Spearman’s correlations<0.11—suggesting that the length of NCPs and CPs does not influence C-end hydrophobicity (Table S3).

We further observed a marked decrease in hydrophobicity along the C-terminal of the longest canonical isoforms, whereas other CPs, NCPs and iORF proteins showed no change in hydrophobicity (Fig. 1C). While these results largely align with those of Kesner et al. (27), they also point to a yet undescribed difference in the C-terminal hydrophobicity between types of canonical proteins (Fig S1).

Confirming selective constraints towards a decreased C-end hydrophobicity, we found significantly higher CTTH scores in shuffled than original in the AUG canonical isoforms (Wilcoxon signed-rank test, see Methods, P < 6.24 x 10^-46^, Fig. 1, Table S2). Conversely, non-AUG isoforms showed comparable CTTH in original and shuffled sequences, which suggests that many of these isoforms might not be translated or have a short half-life. In most NCP datasets, the original sequences shared significantly lower CTTH than shuffled sequences (Fig.1B, Table S2). Nevertheless, the CTTH gap between original and shuffled sequences was much more pronounced in the longest AUG canonical isoforms than in NCPs. Altogether, these results can be explained by selective constraints acting upon a subset of NCPs to reduce CTTH.

In most NCP datasets, CTTH was lower in proteins encoded by uORFs, pseudogenes and internal ORFs (intORFs) than lncRNAs, although this difference was not statistically significant (Fig. 1D, Table S4). Conversely, downstream ORFs showed significantly higher CTTH scores than other classes of NCPs (Table S4). Although CTTH scores were lower in original than shuffled proteins in most cases across datasets, this difference was consistently significant only for uORFs (Figure 1D, Table S4).

### N-end and C-end degrons

Short linear amino acid motifs placed at N- or C-terminal of a protein, known as N-end and C-end degrons, are associated to an increase in degradation propensity (36). We searched CPs and NCPs for 405 N-end and 46 C-end previously characterized degrons that decrease protein stability (37–39) (see Methods). N-end degrons occurred at a significantly higher frequency in NCPs and in short AUG canonical isoforms (41-47%) than in the longest isoforms (pairwise logistic regression with TukeyHSD adjustment, 34%, *P* < 0.003, Fig. 2A, Table S5). Such degrons were also significantly depleted in original compared to shuffled sequences in the longest AUG isoforms (*P* < 0.002). Similarly, C-end degrons were significantly depleted in the longest AUG isoforms than in other CPs and NCPs (pairwise logistic regression with TukeyHSD adjustment, *P* < 9.6 x 10^-7^ in all tests). Both AUG canonical isoforms and most NCP datasets shared fewer C-end degrons than expected given their amino acid composition, particularly in the former (Fig. 2A, Table S6). Across NCP datasets, we observed a significant enrichment of C-end degrons in proteins encoded by uORFs compared to lncRNAs for the Mudge-2022, Chen-2020 and Ji-2015 datasets (pairwise logistic regression with TukeyHSD adjustment, *P* = 0.049, *P* = 0.0008 and *P* = 0.046, respectively; Fig. 2C) and when considering all datasets together (*P* = 2.3 x 10^-8^). Conversely, we found no clear trend for N-end degrons among NCP types (Fig. 2D, Tables S7-8).

**Figure 2.**
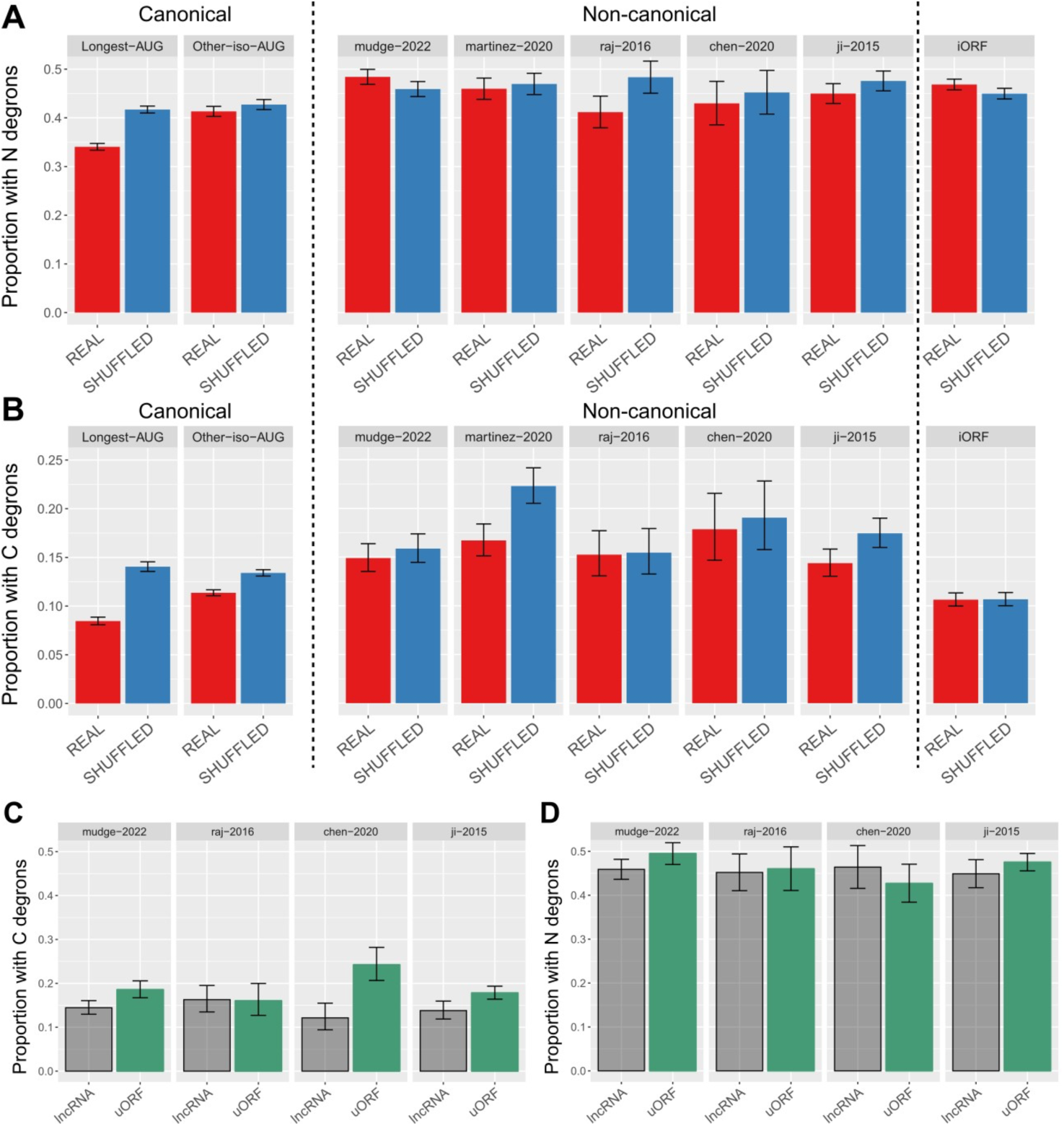
**(A)** Prevalence of 405 N-end degrons in original and shuffled sequences. **(B)** Prevalence of 46 C-end degrons in original and shuffled sequences. **(C)** Prevalence of 46 C-end degrons in peptides encoded by lncRNAs and uORFs in four different datasets. **(D)** Prevalence of 405 N-end degrons in peptides encoded by lncRNAs and uORFs in four different datasets.

### Terminal intrinsic disorder

Terminal intrinsically disordered regions (IDRs) of 30 or more amino acids are associated to protein degradation in mammals and budding yeast (40). We explored if NCPs exhibited different frequencies of IDRs compared to canonical proteins. Only ∼10% of long canonical AUG isoforms possessed either a N-terminal or a C-terminal IDRs, as opposed to the significantly higher 12%-24% of most NCP datasets (Fig. 3A, Fig. S2, Table S9). Intriguingly, the longest canonical AUG proteins had a higher frequency than expected, given the shuffled control sequences, for both N- or C-terminal IDRs (pairwise logistic regression with TukeyHSD adjustment, *P* = 4.6 x 10^-37^ for N-IDRs, *P* = 2.0 x 10^-29^ for C-IDRs), whereas no difference was found in most NCP datasets (Table S9). However, the presence of long IDRs at both termini in the same sequence was significantly lower in canonical AUG isoforms with respect to NCPs (∼2% vs. 5-15%, pairwise logistic regression with TukeyHSD adjustment method, *P* < 1.2 x 10^-13^, Fig. 3B). In most CP but only two NCP datasets, IDRs at both termini were significantly less frequent in original than shuffled sequences, suggesting a strong constraint against this pattern (Table S9). Interestingly, iORFs encoded putative proteins with some of the lowest frequencies of terminal IDRs.

**Figure 3.**
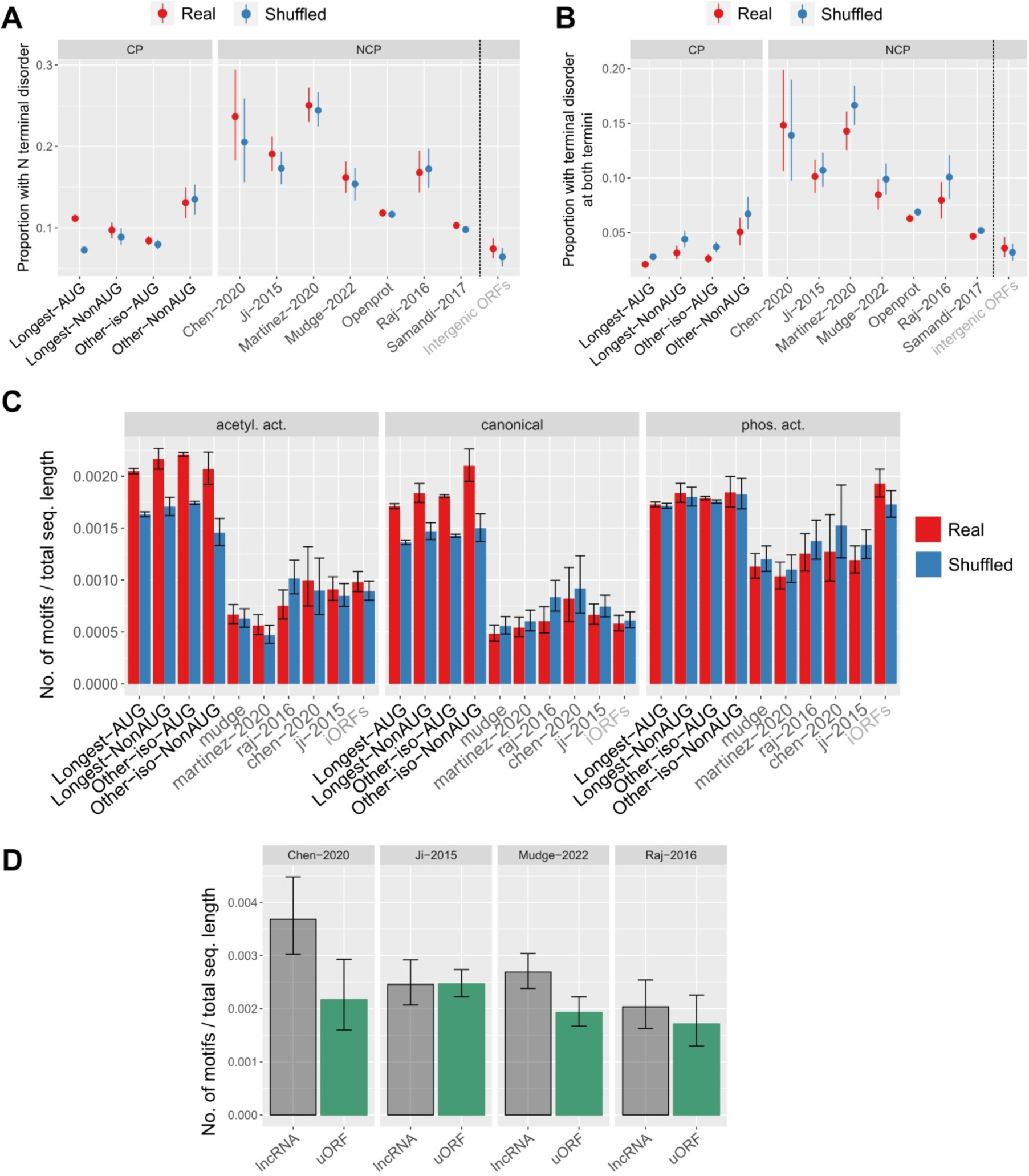
**(A)** Proportion of sequences with N-terminal intrinsically disordered regions. **(B)** Proportion of sequences with C-terminal intrinsically disordered regions. **(C)** Number of KFERQ-like motifs by the total length of each dataset. **(D)** Number of KFERQ-like motifs by the total length of each NCP type dataset.

### Microautophagy-associated KFERQ-like motifs

Chaperone-mediated autophagy (CMA) and endosomal microautophagy (eMI) are responsible for the degradation of specific proteins via lysosomal enzymatic digestion (41). Both these processes involve the recognition of KFERQ-like penta-amino acidic motifs by the chaperone heat shock cognate 71 kDa protein (HSC70). KFERQ-like motifs are either canonical (standard) or formed through posttranslational phosphorylation or acetylation (42). We assessed the possible role of CMA and eMI in noncanonical protein degradation by inferring KFERQ-like motifs occurrence in protein datasets (42). As expected, protein length had a strong correlation with the presence of KFERQ-like motifs (Table S10). Accordingly, 46-74% CPs contained at least one motif compared to 11-18% of NCPs. After normalizing for protein length, the frequency of these motifs remained consistently higher in CPs than NCPs across the three described types of KFERQ-like motifs (canonical, phosphorylation-dependent and acetylation-dependent), and significantly higher in CPs than NCPs overall (*P* = 0 in all tests, Fig. 3C, Table S10). However, KFERQ-like motifs were much more abundant in NCPs than CPs when considering only proteins containing such motifs (Table S10).

For canonical and acetylation-dependent KFERQ-like motifs, original CPs of all categories were significantly enriched compared to shuffled sequences (*P* < 10^-6^ in all tests), whereas this was not the case for NCPs (*P* > 0.05 in all tests, Fig. 3C, Table S10). Intriguingly, the enrichment in the original sequences was entirely absent for the phosphorylation-dependent motifs. Overall, these findings underlie the importance of autophagy-related pathways in the proteostasis of canonical proteins and of a minority of NCPs. We observed a similar pattern across different types of NCPs, with NCPs encoded by lncRNAs sharing higher motif density than those encoded by uORFs in most datasets (*P* < 0.05, Fig. 3D, Table S11).

### Degradations determinants and noncanonical protein evolutionary age

We next tested if degradation propensity in NCPs tend to decrease through time, as observed for CTTH in CPs both in human and mouse (Kesner, et al. 2023). We found no change in both CTTH and terminal IDRs with age in two NCP datasets, including in NCPs more likely to be functional (Figs. S3A-D, Tables S12-3). Similarly, different NCP types showed no change in degradation propensity in relation with their age (Fig. S3E, Table S13). The absence of an evolutionary trend across multiple degradation determinants might simply reflect a biased annotation of increasingly older NCPs as canonical proteins. However, we observed limited to no variation of KFERQ-like motifs with age in CPs as well, suggesting a complex association of gene age with degradation determinants among canonical proteins (Fig. S3F).

### Noncanonical proteins stability and functionality

We next investigated if noncanonical proteins that are more likely to be functional show patterns of stability that align more closely with those observed in canonical proteins. To assess this possibility, we first compared noncanonical sequences with and without an observed phenotype in iPSC and K562 cell lines reported by Chen et al. (10). CTTH scores were significantly lower in proteins with a reported phenotype compared to those with no phenotype (*P* < 0.037), whereas no trend was found for degrons and IDRs (*P* > 0.5, Fig. S4, Table S14). NCPs with a phenotypic effect encoded by lncRNAs and uORFs (except K562 cell lines) also shared lower, but not significant, CTTH values. Original sequences shared lower CTTH than control sequences in proteins with phenotypic effect, but this was significant only for data reported in the K562 line (Table S14). Interestingly, KFERQ-like motifs were consistently less frequent in NCPs with a phenotype (Poisson regression, *P* < 0.001), and in original vs. control sequences (*P* > 0.5), suggesting depletion of CMA motifs in these potentially functional proteins (Table S14).

Second, we analyzed NCPs from a catalog of Ribo-seq ORFs that have been found in one (‘weak’: 4,179 NCPs) or more than one (‘robust’: 3,085 NCPs) studies (21). CTTH was significantly lower in the robust dataset (pairwise logistic regression, *P* = 0.012), whereas all IDRs were significantly higher in the robust dataset (pairwise logistic regression, *P* < 0.05) (Table S15, Fig. S5). N-end degrons, C-end degrons and KFERQ-like motifs occurred more often in the robust datasets but not significantly so (*P* = 0.62, *P* = 0.32 and *P* = 0.65, respectively).

Third, we tested if protein expression correlates negatively with degradation features using the OpenProt database, separating sequences with support by one unique peptide from those supported by at least two unique peptides in mass spectrometry experiments (22). Among OpenProt alternative proteins, which are equivalent to NCPs, CTTH scores were significantly lower (*P* < 0.008) in proteins with at least two supporting peptides (Table S16, Fig. S6A). Conversely, C-end degrons, terminal IDRs and KFERQ-like motifs shared increased frequency in NCPs supported by multiple peptides (*P* < 0.02, Table S16, Fig. S6B-E).

### Degradation signatures and lncRNA localization

LncRNAs occurring only in the nucleus cannot associate to ribosomes and should thus encode putative proteins that freely accumulate degradation determinants compared to lncRNAs primarily found in the cytoplasm. To test this hypothesis, we analyzed data from the LncATLAS database, which contains information on the subcellular localization of lncRNAs in human expressed as Relative Concentration Index, or RCI, between two compartments (43) (see Methods). We identified 3,379 lncRNAs with both nucleus/cytoplasm RCI expression ratio and Ribo-seq data. As expected, ribosome-associated lncRNAs were significantly more likely to localize in the cytoplasm compared to non-translated lncRNAs (Fischer’s exact test, *P* = 0.0001, Table S17). Among lncRNAs with evidence of protein synthesis, those localized in the cytoplasm encoded proteins with significantly lower CTTH than proteins encoded by nucleus-located lncRNA (*P* = 0.0089, Table S17). No other degradation determinants showed significant difference among proteins encoded by cytoplasmic and nuclear lncRNAs (Table S17).

A recently reported lncRNA subcellular localization dataset from human H9 embryonic stem cells (44) was also analyzed applying a cytoplasmic ratio threshold of 0.5 to identify lncRNAs the preferentially localize in the cytoplasm (see Methods). As expected, the cytoplasmic ratio was higher in 1,601 lncRNAs with Ribo-seq detection than 2,102 lncRNAs without a match to Ribo-seq data (Fischer’s exact test, *P* = 0, Table S18). Additionally, the CTTH value was significantly lower in peptides encoded by lncRNAs with cytoplasmic ratio>0.5 and thus more likely to be associated to ribosomes (*P* = 0.0064, Table S18).

### C-terminal tail hydrophobicity and noncanonical tail-anchored proteins

High level of C-end hydrophobicity occurs in canonical tail-anchored (TA) proteins that contain a hydrophobic C-terminal transmembrane domain (TMD) (45, 46). In mammals, TA proteins are recognized by the SGTA/BAG6 checkpoint before being delivered to other mediators of their transmembrane insertion (46, 47), preventing proteasomal degradation despite elevated CTTH (27). Intriguingly, most known human functional NCPs are associated to cellular membranes, suggesting a pathway to functional recruitment of NCPs as novel tail-anchored (TA) proteins (27). We tested this hypothesis more broadly by assessing if high CTTH scores in NCPs are due to an enrichment for TA proteins among noncanonical polypeptides.

The proportion of NCPs with TA proteins-like features was significantly higher compared to AUG canonical proteins, particularly among 83 functional NCP isoforms encoded by 59 ncORFs and reported by Kesner et al. (27) (Fisher’s exact test, *P* < 0.0001, Fig. 4A, Table S19). However, only ∼4-6% of all NCPs might represent functional TA proteins. To determine if NCPs with tail-anchored protein features account for the overall high CTTH in noncanonical polypeptides, we analyzed CTTH in protein datasets after removing predicted TA sequences. CTTH scores remained significantly higher in NCPs compared to canonical longest AUG isoforms (*P* < 4.82 x 10^-5^, Fig. 4B, Table S19).

**Figure 4.**
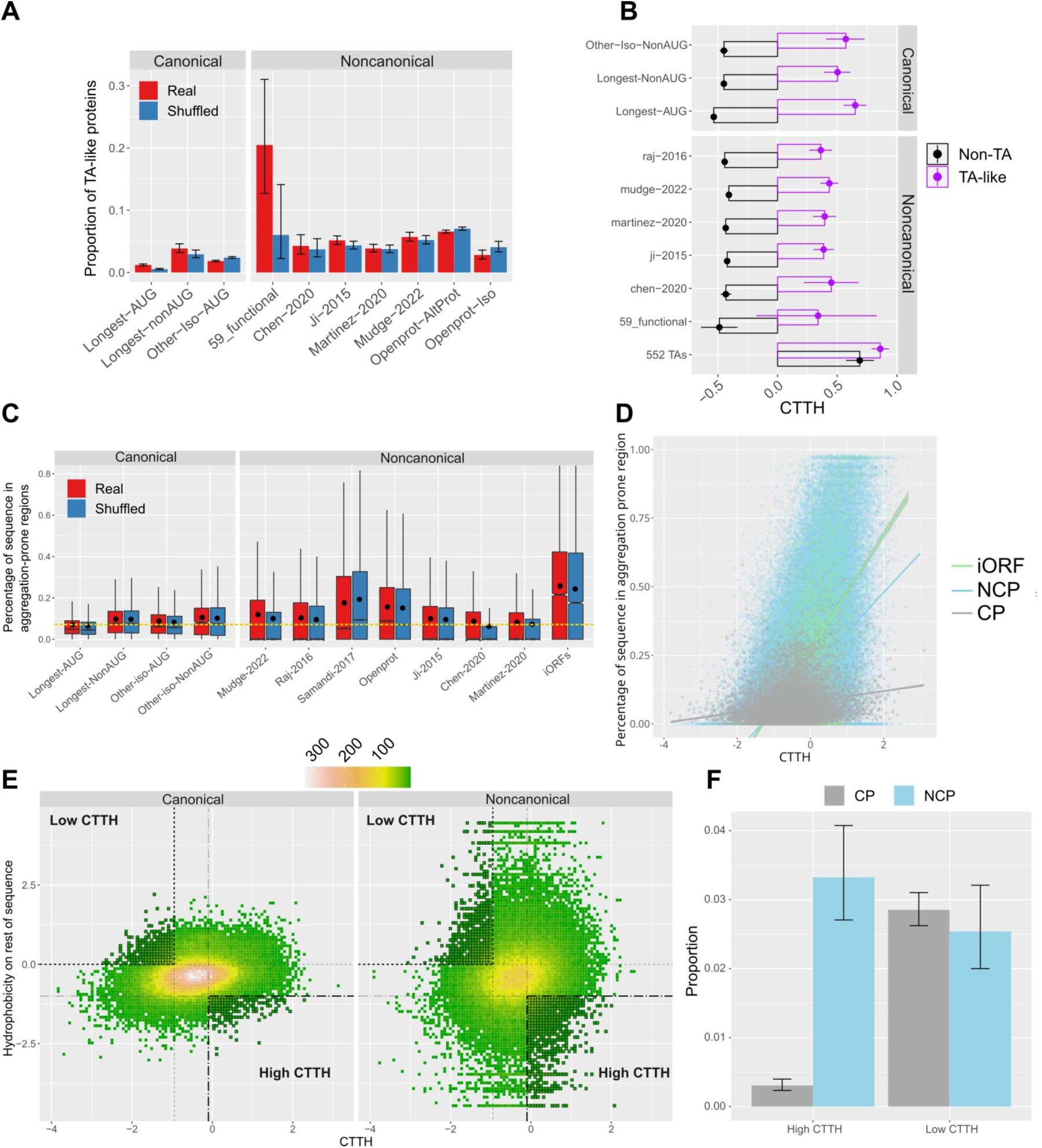
**(A)** Frequency of predicted TA-like proteins in AUG and non-AUG CPs, NCPs and 59 functional NCPs. **(B)** CTTH in TA and non-TA proteins in AUG and non-AUG CPs, NCPs, 59 functional NCPs and 513 human TA proteins. **(C)** Aggregation levels in canonical and noncanonical protein datasets. **(D)** CTTH-Aggregation correlation. Red squares: NCPs. Blue circles: CPs. Black triangles: proteins encoded by intergenic ORFs. **(E)** Distributions of CTTH hydrophobicity in the rest of the protein. Applied cut-offs are drawn as dotted lines. **(F)** Proportion of CPs and NCPs with high and low CTTH values compared to the rest of the protein.

### Degradation determinants and protein aggregation

The formation of toxic aggregates from misfolded proteins is a major cause of cell stress and is associated with a range of diseases (48). Because selection acts to reduce proteins propensity to assemble into large fibrillar structures (49), we hypothesized that degradation determinants could correlate with aggregation propensity. We found that 16-29% of NCP sequences was formed by aggregation-associated amino acids, compared to 5-7% in CPs (*P* < 2.1 x 10^-11^) and 35% in proteins from intergenic ORFs (Fig. 4C; Table S21). No difference between original and shuffled control sequences suggested lack of selection to purge such amino acids.

CTTH was higher in proteins with aggregation propensity than those without, particularly in NCPs. We notice that NCPs exhibited a stronger CTTH-aggregation correlation than CPs (Fig. 4D). However, this correlation increased in shuffled CPs, while it does not change between original and control NCPs (Table S22). Thus, the higher CTTH-aggregation correlation in NCPs might be due to amino acid composition bias rather than selection pressure to decrease stability in more aggregation-prone NCPs. Among other degradation determinants, N-end and C-end degrons showed a weak correlation with aggregation propensity, whereas IDRs were significantly depleted in aggregation-prone proteins, as expected given the inverse correlation between disorder and aggregation (Tables S21-22).

### Patterns of CTTH enrichment and depletion and noncanonical protein function

Stable proteins maintain low C-end hydrophobicity regardless of the overall hydropathy of the whole protein (27). Accordingly, the presence of a combined low CTTH and elevated hydrophobicity in the rest of their sequences is indicative of selective pressure towards high stability. We applied this approach to noncanonical proteins and identified 571/18,583 sequences with highly hydrophilic C-end and a hydrophobic rest of the sequence (see Methods, Fig. 4E-F, Table S20). This is a statistically equivalent proportion than in canonical proteins (552/18,968; Fisher’s exact test, *P* = 0.3798).

We also explored the opposite scenario, whereby natural selection would favor the accumulation of degradation signals, possibly to fine-tune the activity of short-living regulatory noncanonical proteins. We discovered 469 NCPs with both a high CTTH and a strongly hydrophilic rest of the protein, compared to only 54 CPs with the same pattern of hydropathy (Fig. 4E-F, Fisher’s exact test*, P* < 0.00001, Table S20). Thus, the acquisition of a degradation-triggering hydrophobic C-terminal tail may be promoted by natural selection in many more noncanonical than canonical proteins.

## Discussion

The biological impact of noncanonical proteins (NCPs) is controversial due to their potential instability. Here, we tested if NCPs showered lower average stability compared to canonical proteins by assessing the frequency of degradation determinants signatures across hundreds of thousands of human proteins. We investigated three determinants associated with proteasomal degradation: C-terminal tail hydrophobicity, or CCTH (27), N- and C-end degrons (36, 39), and terminal intrinsically disordered regions (40); and amino acid motifs necessary for autophagy-mediated degradation (41, 42). NCPs shared a higher frequency of the three proteasomal-related degradation determinant compared to canonical AUG protein isoforms, underscoring the essential role of multiple surveillance pathways in preventing the expression of potentially harmful proteins. Further supporting these conclusions, we observed diminished degradation signatures in the longest canonical AUG isoforms compared to both non-AUG canonical proteins and NCPs. The longest canonical AUG isoforms also exhibited the strongest selective constrains towards lower CTTH, and reduced frequency of N-end degrons and IDRs at both protein termini.

Noncanonical proteins inferred to be functional are expected to be depleted of degradation features. Accordingly, we found lower CTTH scores and KFREQ-like motif frequencies in noncanonical proteins with a cellular phenotype (10). NCPs expressed in multiple studies (21) and proteomic datasets (22) also shared lower CTTH scores but no difference in KFREQ-like motif occurrence. Thus, experimental phenotypic assays might be more effective in identifying potentially functional noncanonical proteins compared to expression consistency across studies. These findings will inform the ongoing annotation efforts of functional human NCPs (21).

Our results consistently point to CTTH as a reliable predictor of protein stability. The lower CTTH values in cytoplasmic vs. nuclear lncRNAs underscores the importance of this signature in discriminating translated from untranslated ORFs in lncRNAs. Additionally, comparing CTTH scores and hydropathy in the rest of the sequence unveiled hundreds of proteins with either enriched or depleted C-end hydrophobicity, which could result from selective constraints toward a reduced or increased stability, respectively. Notably, sequences with lower than expected stability were significantly more common among NCPs that CPs.

Protein degradation determinants must be accessible to mediators of degradation pathways in order to exert their function, a possible limitation in analyses of stability based on proteins’ primary sequences. For example, KFERQ-like motifs are effective when exposed on the surface of a protein, thus available to autophagy via binding to the chaperone HSC70 (42). However, given that C-terminal tail, degrons and terminal IDRs occur in protein regions that are likely to be exposed (50), most of our findings appear to be robust to possible biases in access to degradation signals.

Most known human functional NCPs are localized on cellular membranes. We confirmed that the majority of these proteins contain a C-terminal transmembrane domain (TMD), a signal present in tail-anchored canonical proteins to allow their appropriate routing to cellular membranes via the SGTA/BAG6 checkpoint (47). TMD are highly hydrophobic and cause the elevated CTTH of TA proteins and membrane-bound NCPs. Since only a small fraction of all NCPs contains a C-terminal TMD, it is unlikely that many NCPs gain functionality by becoming TA proteins.

Because insoluble proteins forming deposits are deleterious in the cellular environment, we hypothesized that the increased instability of NCPs might correlate with their propensity to aggregate. Although NCPs shared stronger aggregation signatures than CPs, we found no evidence that degradation determinants have higher frequency in NCPs more likely to aggregate. Thus, the elevated degradation propensity of NCPs does not seem to result from selection to decrease their potential cellular toxicity due to increased insolubility.

Overall, our study provides an innovative framework to predict degradation propensity in both canonical and noncanonical proteins. We unveiled significant differences in protein stability among >90,000 annotated canonical isoforms that complement and expand on analyses of degradation rates in a few thousands human isoforms (51, 52). Our finding of a consistently lower stability across noncanonical proteins in comparison to canonical isoforms independently substantiate experimental data supporting the defective ribosomal products (DRiPs) hypothesis and demonstrate the power of protein degradation signatures in assessing the impact of the ‘dark proteome’ on human biology.

## Methods

### Datasets

Canonical protein sequences were obtained from the Ensembl v105 database using Biomart. After removing sequences containing the non-standard amino acid “U”, we separated the remaining sequences in two datasets according to the presence/absence of a standard AUG starting codon in the corresponding transcripts. Both AUG and non-AUG datasets were further divided into those including only the longest isoforms of each locus and those containing the rest of the isoforms (Tables S1-S2). Additionally, proteins sharing an identical sequence in the C-terminal 30 amino acids were identified to further remove redundancy in the CTTH analyses.

All NCP datasets were obtained from supplementary tables of the correspondent articles unless differently specified. In the Raj et al. dataset, 122 mCDS and uaCDS with redundant IDs were removed. Genome coordinates of ORFs annotated in hg19 were converted to hg38 coordinates using liftover for the Chen et al. dataset (10) and Raj et al. dataset (9) in order to obtain the corresponding coding regions. Fasta sequences of ORFs were extracted from the hg38 genome assembly using gtf files and the bedtools suite (53). ORFs were translated into protein sequences using the getorf script in the EMBOSS suite (54).

OpenProt contains 48,115 collated entries from 131 ribosome profiling datasets. Entries “Detected with at least two unique peptides” and “Detected with at least one unique peptide” were retrieved from the OpenProt v1.6 dataset (22). Redundant protein sequences between the two datasets were removed.

### Unannotated intergenic genomic ORFs

The hg38 human genome assembly was downloaded together with the gtf files of RefSeq genes (table ncbiRefSeq) and lncRNA and TUCP (Transcripts of Uncertain Coding Potential) from the UCSC Genome Browser. The gtf files of the comprehensive GENECODE V43 dataset was obtained from the GENCODE website https://www.gencodegenes.org/human/. A lncRNAs gtf file (LNCipedia v5.2) was downloaded from https://lncipedia.org/download (55). Unix command lines were used to extract and combine exon sequences from the gtf files into a single file of transcribed exons, which were then merged with bedtools merge. Coordinates of the human genomic regions complementary to transcribed regions were obtained with bedtools subtract on a bed file of hg38. Fasta DNA sequences of these regions were retrieved using bedtools getfasta (53). Sequences containing “N” bases and those <1,000 bp were removed. The coordinates of 1,000 remaining sequences were randomly selected and used to obtain the corresponding genomic DNA from the UCSC Genome Browser. We then used getorf from the Emboss suite (54) to translate the 1,000 random genomic regions.

### Degradation signals analyses

The average hydrophobicity was calculated using the Kyte and Doolittle scale (35), which show high correlation with degradation rates in human proteins (27), using the GRAVY portal https://www.gravy-calculator.de.

Degrons have been experimentally ranked by their contribution to protein instability throughout the differential protein stability index, or ΔPSI. This parameter is calculated as the mean difference in protein stability between polypeptides that contain the degron motif at the N terminus after the initiator methionine (or at the C-end) and polypeptides that contain the same motif at any other internal position (37–39). Degrons with a ΔPSI ≤ −0.4 were retained, as this threshold is well below the ΔPSI value of −0.5 that includes destabilizing motifs (38).

IDR data were obtained using IUPred2a (56) and by parsing the results with in-house python scripts to identify N-terminal and C-terminal IDRs longer than 30 amino acids. Although both terminal and internal IDRs decrease protein stability, we focused only on terminal IDRs because of the length bias of noncanonical proteins. Only non-redundant NPCs longer than 59 amino acids were analyzed.

KFERQ-like motifs were inferred using the ‘motifs from sequence’ option in the KFERQfinder portal (https://rshine.einsteinmed.edu) with default settings to search for canonical motifs, phosphorylation-activated motifs and acetylation-activated motifs (42).

### Protein age

Proteins from the Mudge et al. dataset (21) were divided by Sandmann et al. (57) in four evolutionary ages: human-specific, old world monkey (Catarrhini-specific), primatomorpha (shared by primates and colugos) and conserved across mammals. Information for the noncanonical proteins in Vakirlis et al. (58) included nineteen nodes along the human phylogeny from Vertebrates to human specific. In order to estimate CTTH from at least 4 sequences per age group we collated several age groups. GenTree was used to recover the evolutionary age of human canonical genes (59).

### Known functional microproteins

A list of 64 known functionally characterized microprotein genes was obtained from Kesner et al. (27). Microprotein sequences were retrieved from original papers and the NCBI Protein database. We found some redundancy in the Kesner et al. list, including mitoregulin, which has also been described as MOXI, LEMP and MPM, and BNLN, also named TUNAR. A total of 83 isoform sequences were retained after removing redundant sequences. Cellular phenotypic data from CRISPR assays were obtained from Chen et al.(10).

### LncRNA subcellular localization

Subcellular localization of 6,768 human GENCODE annotated lncRNAs were downloaded from LncATLAS (https://lncatlas.crg.eu). Raw data (2023-05-25_lncATLAS_all_data.csv) were parsed to retain only noncoding sequences. In LncATLAS, the prevalence of a given lncRNA for a subcellular compartment is reported as the Relative Concentration Index, or RCI, corresponding to the log_2_ transformed ratio of the RPKM values from two compartments.

Rows with ‘ratio2’ values, corresponding to the “relative concentration index” (RCI) or log2-transformed ratio of FPKM between the cytoplasm and the nucleus, were retained. When multiple ratio2 values for the same lncRNA were averaged. To retrieve CTTH values for these lncRNAs, we performed Blast analyses against all Ribosomal profiling datasets from Table S1 with calculated CTTH. cDNA sequences of the 6,768 GENCODE lncRNAs were obtained from Ensembl and used in a tBlastn with the following parameters: -ungapped -comp_based_stats F - seg yes -evalue 1e-03 -max_hsps 20 -max_target_seqs 20. Only Blast hits with 100% coverage and identity with at least one RiboSeq sequence were retained. After removing LncATLAS entries with multiple hits, we obtained 3,379 lncRNAs with assigned CTTH values.

Subcellular localization data (nucleus vs. cytoplasm) for lncRNAs in human H9 embryonic stem cells were obtained from Guo et al. (44). The lncRNAs localization was calculated using the cytoplasmic ratio, corresponding to the cytoplasmic FPKMs divided by the sum of cytoplasmic and nuclear FPKMs. We mapped genome coordinates of 4,804 lncRNAs with subcellular information onto the GENCODE v28lift37 lncRNA dataset and found 3,706 unequivocally assigned lncRNA transcripts. tBlastn analyses of these sequences were performed against all Ribosomal profiling datasets from Table S1 with calculated CTTH using the following parameters: -ungapped -comp_based_stats F -seg yes -evalue 1e-03 -max_hsps 20 -max_target_seqs 20. Only Blast hits with 100% coverage and identity with at least one RiboSeq sequence were retained. After removing entries with multiple hits, we obtained 1,601 lncRNAs with assigned CTTH values.

### Prediction of transmembrane domains and tail-anchored (TA) proteins

Transmembrane domains (TMDs) and signal peptides were predicted using the TOPCONS web server using the remote batch script topcons2_wsdl.py (60). The results were parsed using in-house python scripts. We implemented a modified version of the pipeline described by Fry et al. (61) and used the following criteria to define tail-anchored TMDs: absence of a signal peptide and of multiple TMDs, presence of a single TMD no longer than 29 amino acids, a cut-off length for the C-end tail after the TMD of 29 amino acids. Non-TA proteins with TMDs were identified as any remaining sequence with a TMD. Our approach was robust to false positives, as we identified ∼60% of TA proteins described by Fry et al. (61), likely due to the elevated sensitivity to signal peptide detection in TOPCONS, which are considered incompatible with TA proteins. Our datasets also included 83 isoforms from a list of 59 non-redundant functionally characterized microproteins reported in Kesner et al. (27), most of which have been found to localize on cellular membranes and thus are expected to possess transmembrane domains.

### Hydropathy and functional noncanonical proteins

We calculated the Kyte-Doolittle index for the last 30 amino acids, i.e., the CTTH score, and for the rest of the sequence in noncanonical proteins longer than 59 amino acids. Proteins with CTTH below the lowest quartile in AUG canonical longest isoforms (CTTH < −0.95) and a hydrophobic (positive) Kyte-Doolittle index in the rest of their sequence were retrieved as possible examples of proteins selected to reduce degradation propensity. To identify cases that represented the opposite pattern of selection to increase instability, we used as thresholds a CTTH above the highest quartile of AUG canonical proteins of −0.107 (after removing proteins with a transmembrane domain at the C-end as predicted using TOPCONS (60)) and a highly hydrophilic rest of the protein (Kyte-Doolittle index<-1).

A total of 405 experimentally characterized human N-degrons falling below this threshold were obtained from Timms and co-authors (38). Forty-six C-degrons with ΔPSI ≤ −0.4 were obtained from Koren et al. (37). Analyses were conducted both with the full list of 405 N-degrons and 46 C-degrons as well as with the top 20 motifs in each group according to their ΔPSI. Sequences with identical 30 amino acids at their N-end and C-end were removed from the N-degron and C-degron analyses, respectively. Sequences with identical 30 amino acids at the C-end were also removed from the CTTH results.

### Statistical analyses

All statistical analyses and data representations were performed using the Rv4.3 environment (62). All *p-*values for comparisons between datasets were calculated using the Wilcoxon rank sum test, except where differently indicated. A continuity correction and Bonferroni *P* value adjustment were applied when multiple datasets comparisons were performed. All *p*-values for original vs. shuffled sequences were calculated using the Wilcoxon signed-rank test. Confidence intervals of proportions were calculated with R function prop.test.

## Acknowledgments

We thank Dr. Dimitris Nakos for fruitful discussions and feedback regarding the manuscript. This study was supported by the National Institute of Food and Agriculture, U.S. Department of Agriculture, under award number TEX0-1-9599, the Texas A&M AgriLife Research, the Texas A&M Forest Service, and the Hellenic Foundation for Research and Innovation (H.F.R.I.) under the “3rd Call for H.F.R.I. Research Projects to support Post-Doctoral Researchers” to N.V. (Project Number:7330).

## Supporting Figures

**Supporting Figure 1.**
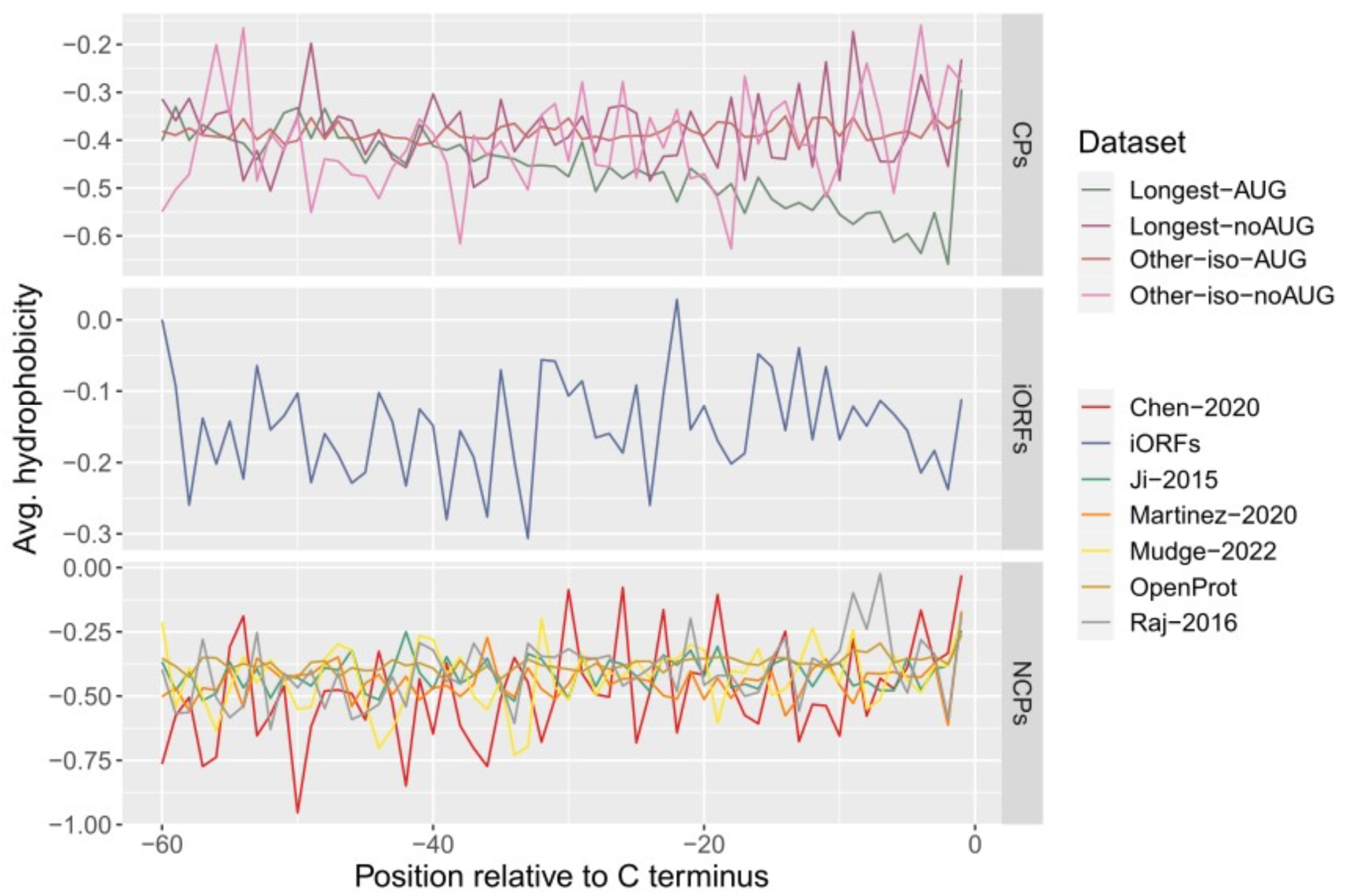
Average hydrophobicity (Kyte-Doolittle scale) in the last 60 amino acids of CPs, NCPs and iORFs.

**Supporting Figure 2.**
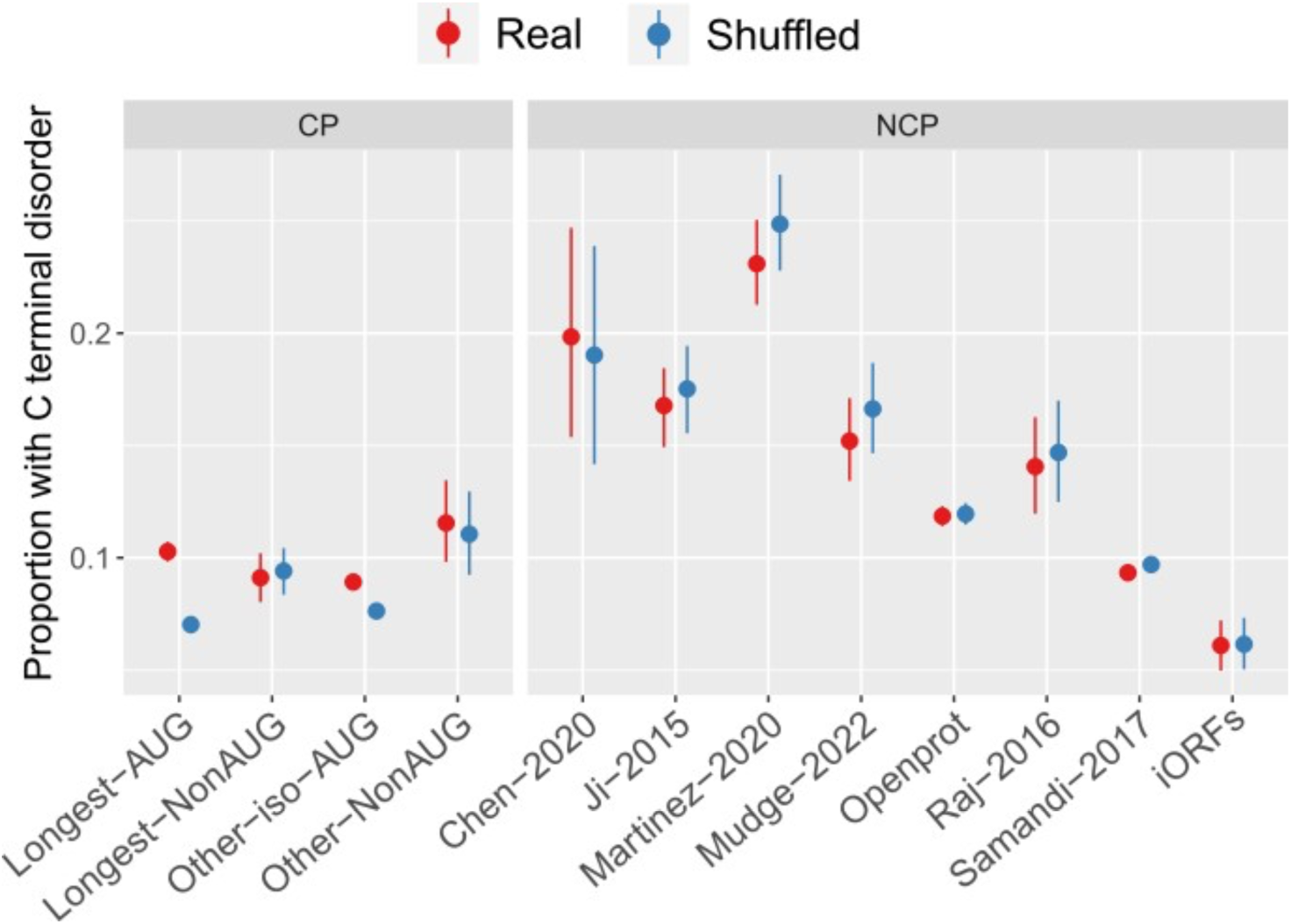
Proportion of sequences with C-terminal intrinsically disordered regions.

**Supporting Figure 3.**
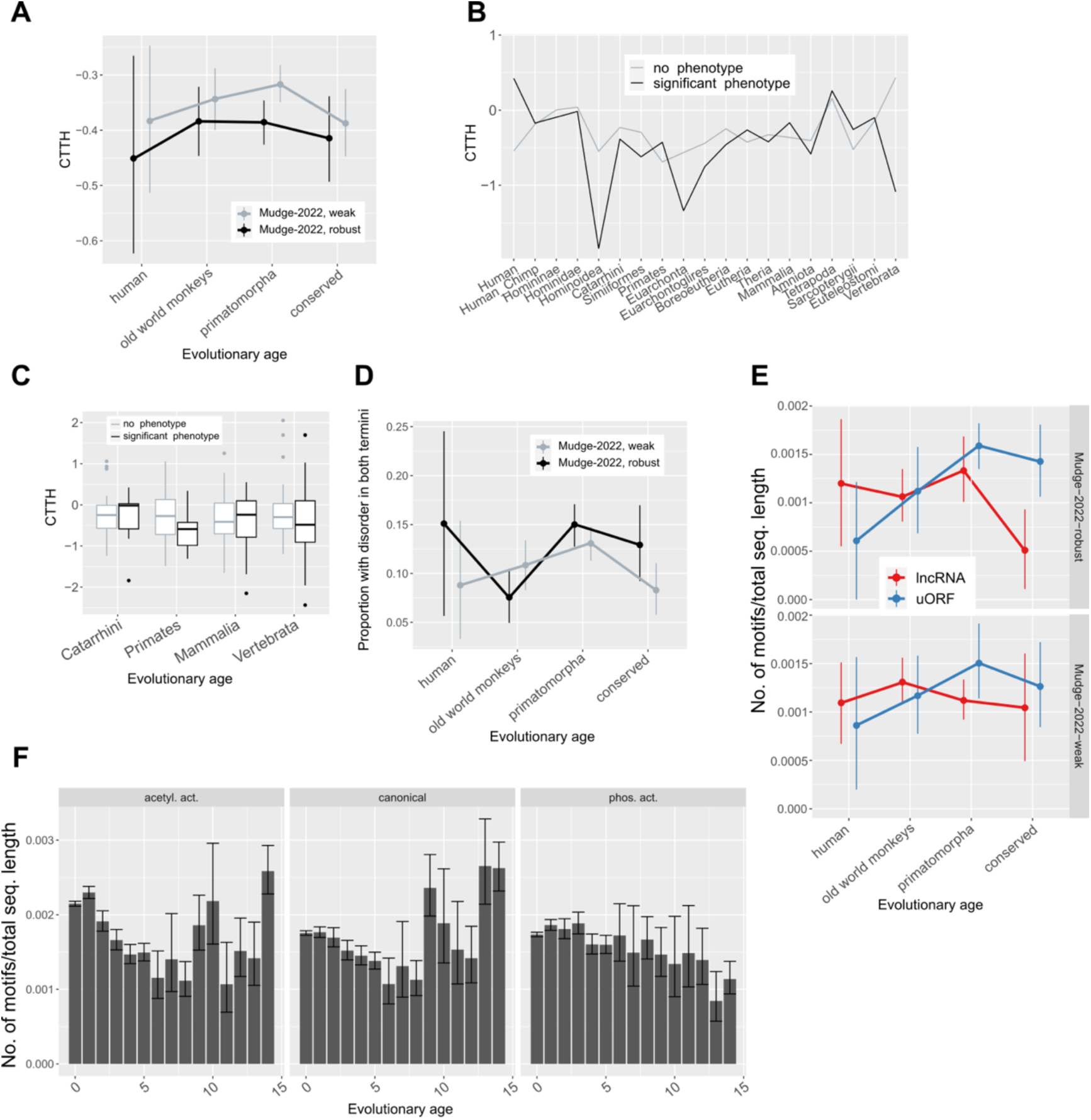
**(A)** Average CTTH in noncanonical proteins with increasing age from left to right as assessed by Sandmann et al. (57) on data from Mudge et al. (21) (weak and robust datasets). **(B)** Average CTTH in NCPs with increasing age from left to right as assessed by Vakirlis et al. (58) on data from (10). Microproteins with significant phenotypes and no phenotypes as measured by Chen et al. are shown separately. **(C)** Same as B but on four broader groups of evolutionary age. **(D)** Frequency of KFERQ-like motifs in NCP types from the robust and weak datasets from Mudge et al. (2022)., with increasing age from left to right as assessed by Sandmann et al. (57). **(E)** Frequency of IDRs on both termini in NCPs assessed by Sandmann et al. (57) on the robust and weak datasets from Mudge et al. (21). **(F)** Frequency of KFERQ-like motifs of different types, in longest canonical isoforms of annotated proteins, with decreasing evolutionary age as assessed by Shao et al. (2019).

**Supporting Figure 4.**
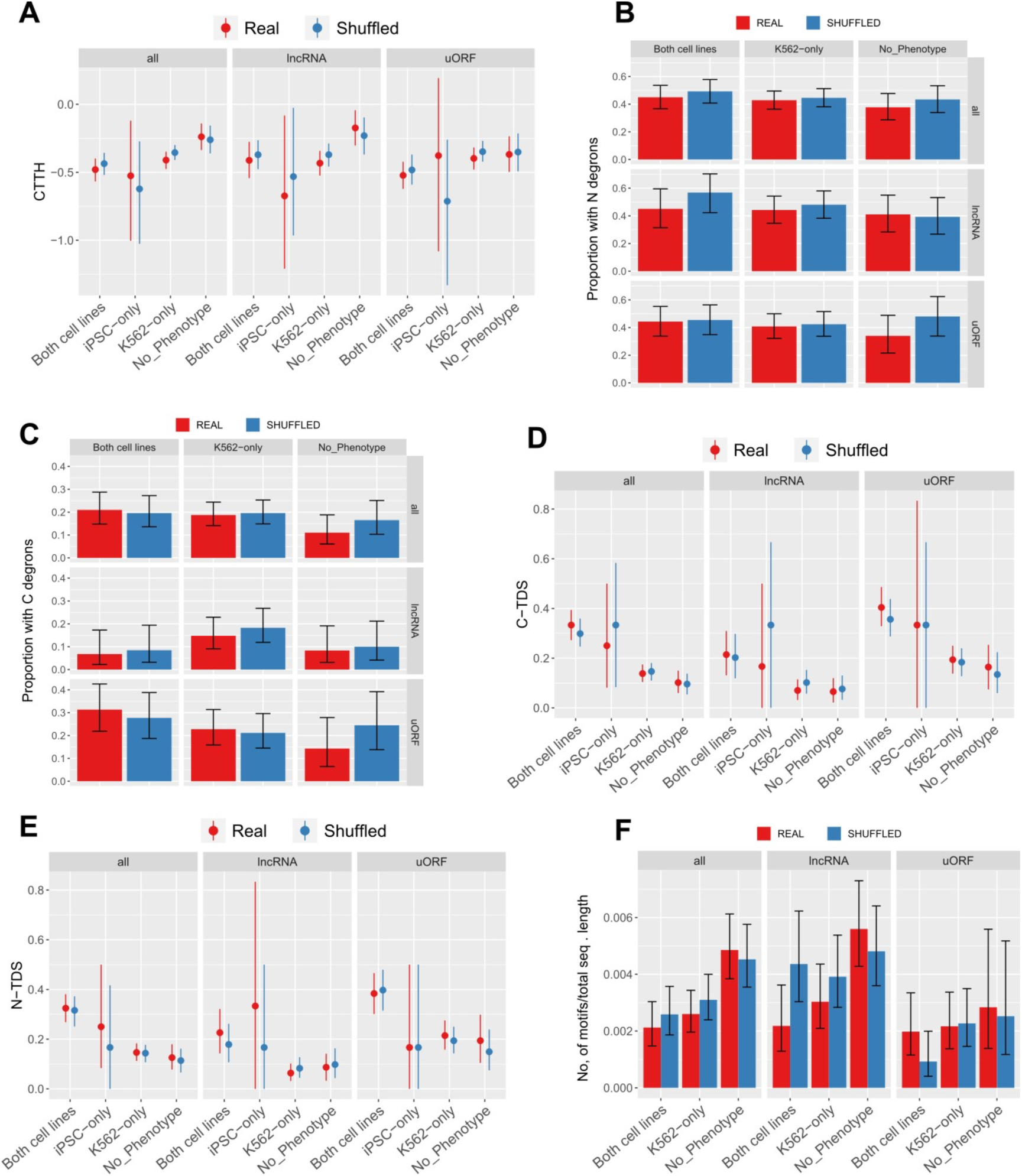
Degradation determinants in noncanonical proteins tested in K562 and iPSC cell lines. **(A)** CTTH. **(B-C)** Degrons. **(D-F)** IDRs. **(G)** KFERQ-like motifs frequency. K562: proteins with phenotype in K562 cells. Both: proteins with phenotype in K562 and iPSC cells. NoPh: proteins with no phenotype.

**Supporting Figure 5.**
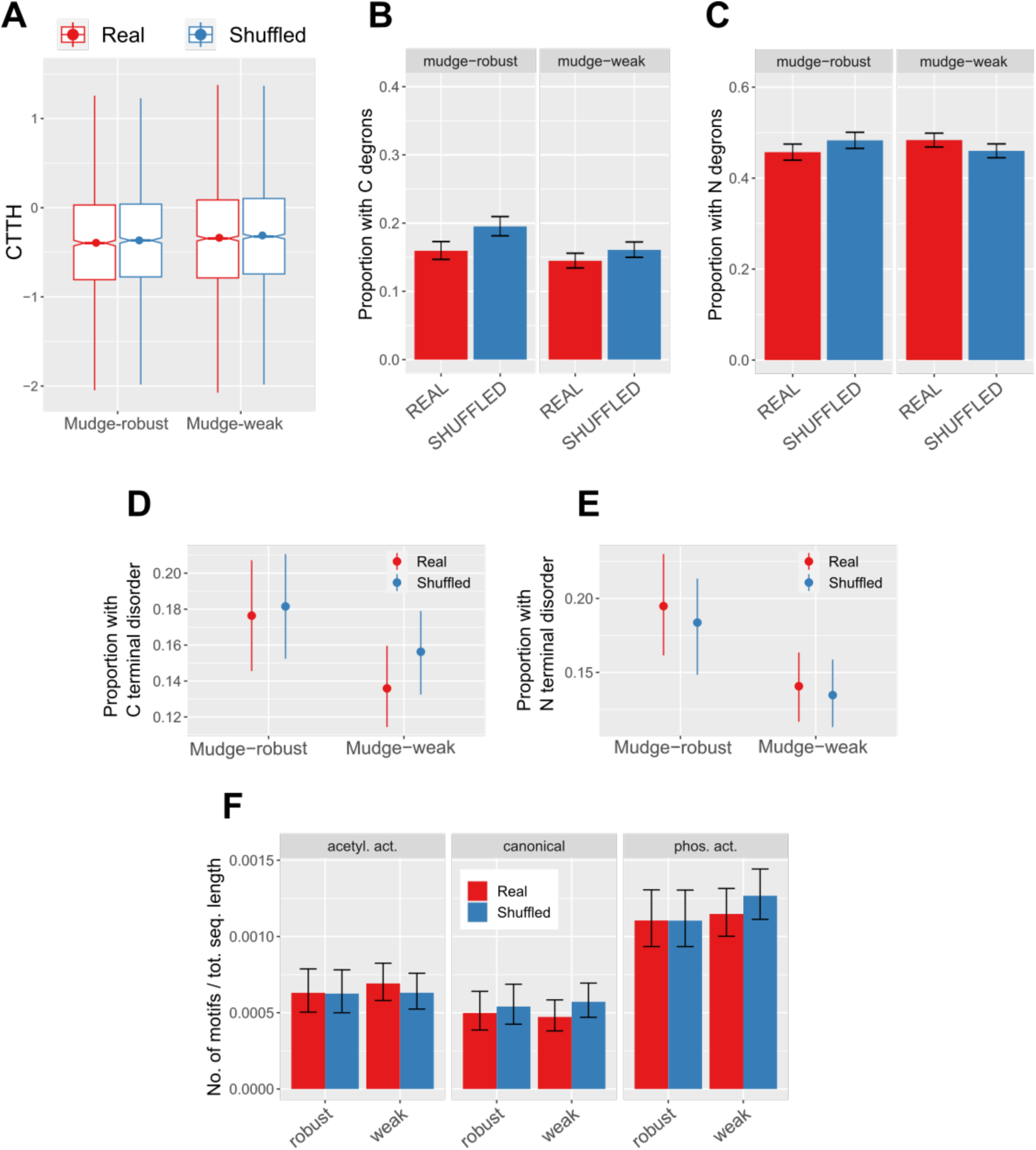
Degradation determinants in noncanonical proteins expressed in one study (weak) or multiple studies (robust). **(A)** CTTH. **(B-C)** Degrons **(D-F)** IDRs. **(G)** KFERQ-like motifs frequency.

**Supporting Figure 6.**
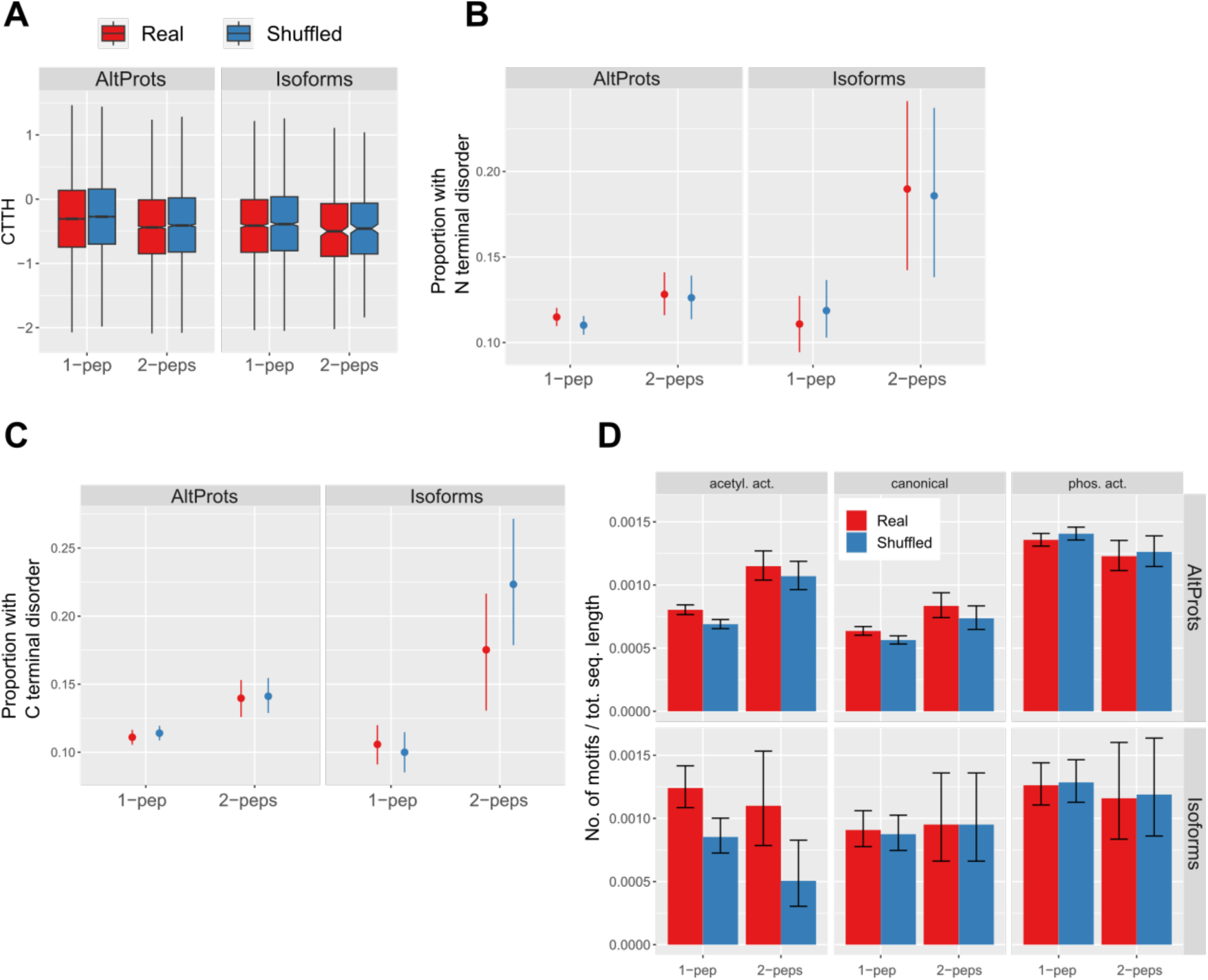
Degradation determinants in OpenProt noncanonical proteins with varying degree of proteomic support. (**A)** CTTH. **(B-C)** IDRs. **(E)** KFERQ-like motifs frequency.

## Notes

https://figshare.com/s/bb5a3f24133ad5540688

### Competing Interest Statement

The authors have declared no competing interest.

